# Mass measurements of polyploid lymphocytes reveal that growth is not size limited but depends strongly on cell cycle

**DOI:** 10.1101/2019.12.17.879080

**Authors:** Luye Mu, Joon Ho Kang, Selim Olcum, Kristofor R. Payer, Nicholas L. Calistri, Robert J. Kimmerling, Scott R. Manalis, Teemu P. Miettinen

## Abstract

Cell size is believed to influence cell growth and metabolism. Consistently, several studies have revealed that large cells have lower mass accumulation rates per unit mass (i.e. growth efficiency) than intermediate sized cells in the same population. Size-dependent growth is commonly attributed to transport limitations, such as increased diffusion timescales and decreased surface-to-volume ratio. However, separating cell size and cell cycle dependent growth is challenging. To decouple and quantify cell size and cell cycle dependent growth effects we monitor growth efficiency of freely proliferating and cycling polyploid mouse lymphocytes with high resolution. To achieve this, we develop large-channel suspended microchannel resonators that allow us to monitor mass of single cells ranging from 40 pg (small diploid lymphocyte) to over 4000 pg, with a resolution ranging from ~1% to ~0.05%. We find that mass increases exponentially with respect to time in early cell cycle but transitions to linear dependence during late S and G2 stages. This growth behavior repeats with every endomitotic cycle as cells grow in to polyploidy. Overall, growth efficiency changes 29% due to cell cycle. In contrast, growth efficiency did not change due to cell size over a 100-fold increase in cell mass during polyploidization. Consistently, growth efficiency remained constant when cell cycle was arrested in G2. Thus, cell cycle is a primary determinant of growth efficiency and increasing cell size does not impose transport limitations that decrease growth efficiency in cultured mammalian cells.

**Significance statement:** Cell size is believed to influence cell behavior through limited transport efficiency in larger cells, which could decrease the growth rate of large cells. However, this has not been experimentally investigated due to a lack of non-invasive, high-precision growth quantification methods suitable for measuring large cells. Here, we have engineered large versions of microfluidic mass sensors called suspended microchannel resonators in order to study the growth of single mammalian cells that range 100-fold in mass. This revealed that the absolute size of a cell does not impose strict transport or other limitations that would inhibit growth. In contrast to cell size, however, cell cycle has a relatively large influence on growth and our measurements allow us to decouple and quantify the growth effects caused by cell cycle and cell size.

## Introduction

The extent to which cell cycle and cell size affect cell growth efficiency (growth rate per unit mass) is not known. In cultured and proliferating animal cells, mass increases exponentially with time, except in the largest cells which display decreased growth efficiency and proliferation rates (1–4). One explanation for the decreased growth in largest cells is that when cells grow beyond a certain size their growth becomes constrained by transport limitations (5–14). Most notably, larger cells have longer diffusion distances and lower surface-to-volume ratios, both of which could reduce the maximal rate at which large cells can transfer metabolites and information. Importantly, such transport limitations can exist even when cellular components scale isometrically with cell size. In a developmental setting, growth-influencing transport limitations could have a major impact on cell physiology, possibly explaining why most fast growing and proliferating cell types are small (<20 μm in diameter) (5, 6, 8). Transport limitations are also considered to result in allometric scaling of metabolism, a phenomenon where larger animals display lower metabolic and growth rates (10, 11). However, whether increasing cell size fundamentally imposes transport limitations that result in decreased growth efficiency is not known.

Alternatively, the non-linear correlation between cell mass and growth efficiency could reflect cell cycle dependent growth, where each specific cell cycle stage has differential growth signaling and metabolism. This growth regulation can be entirely independent of cell size or can be coupled to size-dependent titration/dilution effects, where the concentration of cellular components is lowered as cells grow larger. Such dilution effects often depend on DNA content and, consequently, the dilution effects should be most prominent when cells grow during a cell cycle arrest (14–18). In support of cell cycle dependent growth, cell cycle regulators are known to influence protein synthesis machinery (19–22), and growth rates in G1 have been shown to depend on cell size (23, 24), presumably due to dilution effects. However, as cell cycle stage changes with cell size in most proliferating cell types, cell size and cell cycle effects must be decoupled to understand their individual contributions to cell growth.

To quantify the extent of cell size dependent growth, one would need to examine cells of vastly different sizes. Cultured cells maintain size homeostasis and display little size variability. Yet, cell size can increase significantly when cells undergo repeated cell cycles in the absence of cell division (polyploidization). Polyploidization and the associated cellular hypertrophy is normal and critical in many tissues during development (5, 12, 25, 26), and also commonly observed in cancers (25, 27). Although the physiological importance of polyploidy is well established, method limitations have prevented high-resolution single-cell measurements of growth in large polyploid cells. Several methods, including quantitative phase microscopy (3, 28), fluorescence exclusion microscopy (23, 29) and suspended microchannel resonators (SMRs) reported thus far (2, 30), are capable of non-invasively quantifying single-cell growth rates of small cells (diameter range from <5 μm to 15 μm in spherical cells). However, for the large cell sizes observed in polyploid cells, these techniques become imprecise or even infeasible, depending on the method. Here, we expand the analytical range of SMRs by engineering large-channel versions of the devices. We then use the large-channel SMRs together with previously published small-channel SMRs to monitor the growth of vastly different sized single cells (ranging from ~7 to ~32 μm in diameter) and to quantify the extent to which cell size and cell cycle influence growth.

## Results and discussion

The SMR is a microfluidic mass measurement device where a cell is flown through a vibrating cantilever and the change in the cantilever’s vibration frequency is used to quantify the buoyant mass of the cell. To overcome previous size-range limitations, we developed large-channel SMRs, which have 60 × 60 μm microfluidic channel inside the vibrating cantilever (Fig. 1A). These large-channel devices operate in the first vibration mode and utilize a new image-based hydrodynamic trapping approach to repeatedly measure the buoyant mass of a single particle/cell *(SI Appendix,* Fig. S1, Materials and Methods). The image-based hydrodynamic trapping provided additional stability for long-term mass monitoring by allowing us to maintain a cell or a bead in a specified region within the microfluidic channels between measurements. Using polystyrene beads, we quantified each large-channel SMR mass measurement to have a resolution (standard deviation) ranging between 0.24 and 1.25 pg for particles ranging from 10 to 50 μm in diameter, respectively (Fig. 1B; see *SI Appendix,* Fig. S1 & Materials and Methods for full details). This corresponds to a measurement coefficient of variation range from 1.1 % to 0.05 %, respectively. When monitoring single-cell growth, we were able to carry out mass measurement every ~30 s without affecting cell viability, allowing us to average multiple mass measurements when monitoring mass changes that take place over longer time periods (Fig. 1C).

**Fig. 1.**
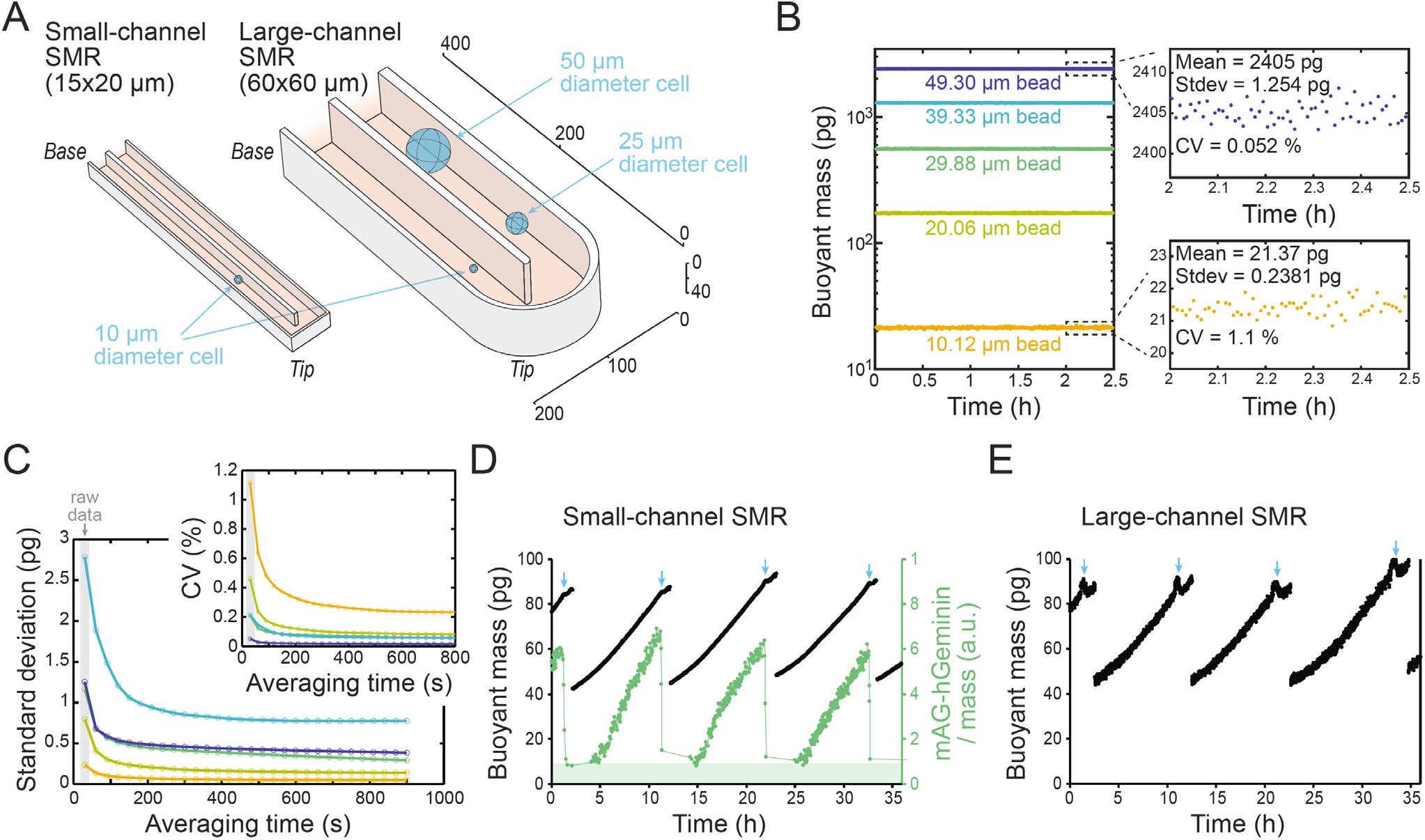
Large-channel SMR enables buoyant mass monitoring across large size ranges with high resolution. **(A)** In scale schematic of the small and large-channel SMR cantilevers. The measurement principle of SMRs is to flow a cell through a vibrating cantilever while monitoring changes in the vibration frequency, which are directly proportional to the buoyant mass of the cell. Note that small-channel SMRs are equipped with fluorescence detection system, whereas large-channel SMRs use image-based cell trapping. The scale bars on the right are in μm. **(B)** Quantification of large-channel SMR resolution based on repeated mass measurements of single polystyrene beads of different sizes (diameters). Insets display zoom-in views of the 10.12 and 49.30 μm bead data along with measurement mean, standard deviation (stdev) and coefficient of variation (CV). **(C)** Large-channel SMR mass measurement resolution as a function of averaging time (moving average filter length reflecting temporal resolution) over multiple measurements for different sized polystyrene beads. Color coding is the same as in panel (B). The measurement interval is ~30 s, and the first data point under a grey background reflects individual measurements without any averaging. Inset displays the resolution as CV. **(D,E)** Example mass trace of control L1210 FUCCI cell growing through multiple divisions in (D) small-channel SMR (N=9 independent experiments) and in (E) large-channel SMR. Data represents individual mass measurements without any averaging. At each division, one daughter cell is discarded. The mAG-Geminin signal (green) detection was only carried out in small-channel devices, and its increase indicates G1/S transition and loss indicates metaphase/anaphase transition (blue arrows).

To validate that the large-channel SMRs provide data comparable to previous 15 × 20 μm SMRs (from here on referred to as small-channel SMRs), we measured single-cell buoyant mass accumulation rate (from here on referred to as growth rate) of mouse lymphocyte L1210 cells expressing the mAG-hGeminin cell cycle reporter (FUCCI). The interphase L1210 cell growth rates obtained from small and large-channel SMRs were similar (Fig. 1D and E). It is known that growth rate, cell density and cell stiffness display dynamic changes in mitosis (19, 31, 32). As these changes are unlikely to reflect cell size-dependent effects, we have excluded mitosis from all future analyses. While the small-channel SMR has better measurement resolution than the large-channel SMR when measuring normal sized L1210 cells (stdev of 0.026 pg and 0.24 pg for a 10 μm diameter bead, respectively) (19), the large-channel SMR increases the maximum spherical cell volume that can be measured 64-fold. Importantly, the large-channel SMR is also able to monitor growth of a single cell over multiple cell cycles (randomly following one of the daughter cells following each division, Fig. 1E), which is previously achieved by only a few cell size measurement methods (33).

We first studied the size-dependency of cell growth by monitoring unperturbed L1210 cells using the small-channel SMRs. Our data revealed that when cells are small (G1 and early S-stage cells), growth rate increases linearly with size (as cell cycle proceeds), indicative of exponential growth (Fig. 2A). However, growth rate plateaus in larger cells (late S-stage and G2 cells), indicating a transition to linear growth. Consequently, the intermediate sized cells (S-stage cells) displayed the highest growth efficiency (Fig. 2B). Large-channel SMRs provided similar data. These results are consistent with previous findings (1–3, 23) that cell size and/or cell cycle have a major effect on cell growth efficiency.

**Fig. 2.**
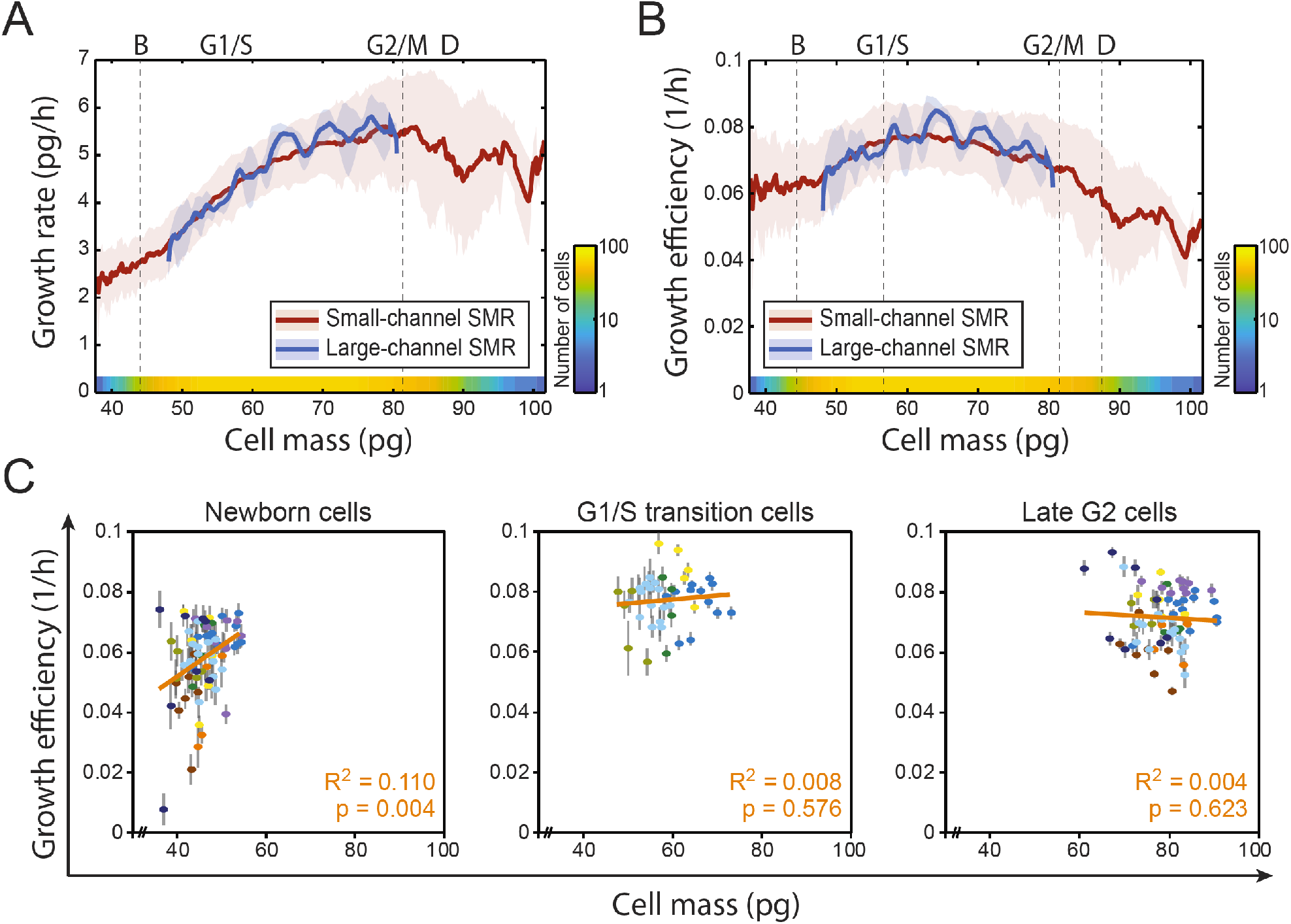
Growth efficiency of unperturbed lymphocytes does not correlate with cell size when examining specific cell cycle stages. **(A,B)** The growth rate (A) and growth efficiency (B) of L1210 FUCCI cells as a function of mass as obtained using small-channel SMR (red trace, number of cells for each size analyzed is indicated with a color gradient at the bottom, N=9 independent experiments) and large-channel SMR (blue trace, n=3). The line and shaded area indicate mean ± stdev. Average newborn size (B), G1/S transition size, mitotic entry size (G2/M) and division size (D) are indicated with dashed vertical lines. **(C)** Correlations between mass and growth efficiency at the beginning of G1, at G1/S transition and at the end of G2 (n=72 cells from 9 independent experiments for newborn and late G2 cells; n=41 cells from 5 independent experiments for G1/S cells). Data from each experiment is represented in its own color. Each cell (dot) is plotted with error bars (measurement error as stdev; error bars for cell mass are smaller than the dots). Linear fits, Pearson correlations (R^2^) and p-values for the correlations (2-tailed test of significance) are shown in orange.

Next, to examine cell cycle stage independent growth effects, we measured growth efficiency specifically in newborn G1 cells, in cells at G1/S transition and in cells at late G2. This revealed little to no correlation between growth efficiency and cell mass (Fig. 2C). Considering our measurement resolution (19), the lack of correlation is unlikely to be caused by noise in our measurement. Thus, these results suggest that cell size does not have a major influence on L1210 cell growth efficiency when examining changes over small size ranges.

To examine size-dependent growth independently of cell cycle effects and over a large size range, we turned to polyploid model systems. If the declining growth efficiency observed in the largest unperturbed cells (Fig. 2B) is due to transport limitations caused by increases in cell size, then increasing size further by induction of polyploidy should result in further declining growth rates (Fig. 3A). However, if the non-linear correlation between cell size and growth efficiency observed in control cells reflects cell cycle dependent growth, then the oscillating growth efficiency should repeat with every successive cell cycle in the polyploid cells independently of cell size. To test our hypothesis, we induced polyploidy in L1210 cells using 50 nM Barasertib (also known as AZD1152-HQPA), a selective inhibitor of Aurora B, which is critical for cytokinesis (34, 35). This resulted in several endomitotic cycles where ploidy increased from 2N up to 128N (Fig. 3B) with corresponding increases in cellular hypertrophy (Fig. 3C and D), suggesting that DNA-to-cell size ratio remained comparable to control cells. Importantly, the cells remained spherical with a single, multilobed nucleus (Fig. 3C). Prolonged drug treatments also resulted in cell death, which manifested in mass measurements as sudden transition to zero or negative growth (*SI Appendix*, Fig. S2A-C). These data were excluded from our analysis (Materials and Methods).

**Fig. 3.**
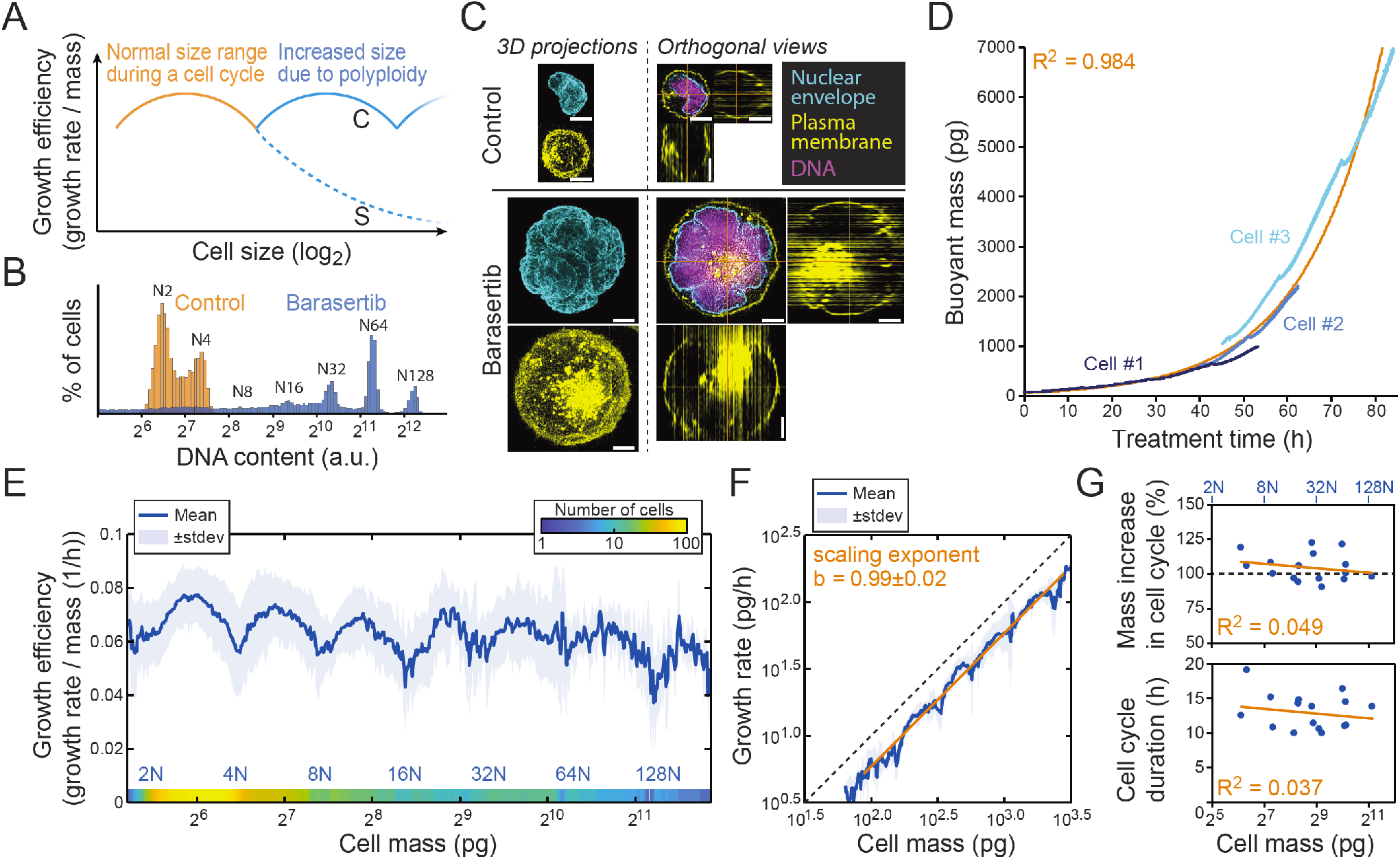
Monitoring polyploid cell growth over a 100-fold size range reveals that growth is not size limited. **(A)** Hypothesis of cell growth regulation as cells increase size through endomitotic cycles. In the size range observed during a normal cell cycle (orange) cells display non-linear size scaling of growth efficiency. When cells grow in to polyploidy, cell size dependent (S, dashed blue line) and cell cycle dependent (C, solid blue line) growth should result in different growth behaviors. **(B)** DNA histogram of untreated (orange) and 80h 50 nM Barasertib treated (blue) L1210 FUCCI cells (n=3 independent cultures). **(C)** Examples of control (top) and 50 nM Barasertib treated (bottom) L1210 FUCCI cell morphologies based on nuclear envelope (cyan), plasma membrane (yellow) and DNA (magenta) imaging (N=2 independent experiments each with >10 fields of view). Left side displays 3D projections and right side displays a single z-slice with orthogonal views. Scale bars denote 5 μm. **(D)** Three example buoyant mass traces from Barasertib treated L1210 FUCCI cells obtained using the large-channel SMR (N=31 independent experiments). Exponential fit to the three examples and Pearson correlation (R^2^) are displayed in orange. **(E)** The combined growth efficiency of control, Barasertib and H-1152 treated L1210 FUCCI cells across a large mass range as measured with both small and large-channel SMRs. Number of cells at each mass is indicated with a color gradient at the bottom (N=76 independent experiments). Estimated ploidy level is displayed on bottom in blue. See also SI Appendix, Fig. S2E and F. **(F)** Growth rate as a function of mass on a log_10_-log_10_ plot when utilizing only Barasertib treated L1210 data collected using the large-channel SMR. Linear fit and scaling exponent *b* (mean ± s.e.m.) are displayed in orange. Perfect isometric scaling (*b* = 1) is illustrated with dashed black line. **(G)** Correlation between % mass increase during a cell cycle *(top)* or cell cycle duration (*bottom*) and cell mass at the beginning of the cell cycle. Data represents Barasertib treated L1210 cells analyzed using the large-channel SMR (N=11 independent experiments with 16 cell cycles). Dashed black line in top panel represents a perfect mass doubling in each endomitotic cycle. Approximate ploidy level at the start of each cell cycle is displayed on top in blue. Linear fits and Pearson correlations (R^2^) are displayed in orange.

When examining growth over larger size scales using the polyploid cells, mass increased exponentially over time (Fig. 3D and *SI Appendix,* Fig. S2D). Remarkably, the non-linear growth efficiency behavior that was observed in control cells (Fig. 2B) repeated in every successive cell cycle during polyploidization independently of cell size (Fig. 3E). This non-linear growth behavior cannot be explained by the DNA-to-cell size ratio alone, as growth efficiency decreased towards the end of each cell cycle, but started to increase immediately following endomitosis before the subsequent S-stage. Furthermore, the low growth efficiency in newborn G1 cells (Fig. 2B and C), cannot reflect too small absolute size, as polyploid G1 cells display similarly low growth efficiency. To validate that the observed growth behavior cannot be attributed to drug specific effects, we induced polyploidy using an alternative cytokinesis inhibitor, 10 μM H-1152, which targets the Rho-kinase (ROCK) (36). This resulted in similar growth behavior as Aurora B inhibition *(SI Appendix,* Fig. S2E).

Overall, we quantified growth efficiency for L1210 lymphocytes over an approximately 100-fold mass range spanning from 40 pg to 4000 pg. In spherical L1210 cells this corresponds to a diameter range from <7 μm to >32 μm resulting in estimated 4.5-fold reduction in surface-to-volume scaling. This size range covers most proliferating cell types in the human body. Unlike cells *in vivo,* cultured cells are constantly selected for the highest growth rate, allowing us to assume that the measured growth rates reflect maximal growth rates possible for the cells. Size scaling typically follows a power law *Y = aM*^ь^, where *Y* is the observable biological feature, *a* is a normalization constant, *M* is the mass of the organisms (or a cell), and *b* is the scaling exponent which typically has values close to ¾ when studying metabolic rate (10, 11). We observed a minor decrease in growth efficiency in largest cells when plotting data obtained across multiple measurement systems and conditions (Fig. 3E). We therefore quantified size-dependent growth and the allometric scaling exponent from our growth rate data using only Barasertib-treated cells monitored with the large-channel SMR *(SI Appendix,* Fig. S2F). After accounting for the oscillating cell cycle dependent growth (Materials and Methods), the L1210 cell growth rates display an isometric scaling exponent of 0.99 ± 0.02 (mean ± s.e.m.) (Fig. 3F), consistent with previous predictions for cells *in vitro* (37). This corresponds to each doubling of cell mass changing growth efficiency by −0.1 ± 1.1% (mean ± s.e.m.).

In contrast to cell size, cell cycle displays a strong influence over cell growth efficiency. To validate that cell cycle progression causes the oscillating growth behavior within each cell cycle, we arrested L1210 cells to G2 stage with 2 μM RO-3306, an inhibitor of cyclin-dependent kinase 1 (CDK1) (38) (*SI Appendix,* Fig. S3A and B). Prolonged RO-3306 treatment resulted in cell death, and to avoid this toxicity, we only analyzed growth for the first 40 pg increase (corresponding to a typical mass increase in a cell cycle) from the normal mitotic size. This revealed that the decrease in growth efficiency that was observed in large control cells stopped as cells were arrested in G2 and the growth efficiency remained constant for G2 arrested cells even as their sizes increased *(SI Appendix,* Fig. S3C and D). Thus, as suggested by previous work in budding yeast (39), our results show that cell cycle has a major influence on mammalian cell growth efficiency. We quantified this cell cycle dependent growth to be 29 ± 3% (mean ± s.e.m.) of the average growth efficiency in untreated L1210 cells. In addition, the steady growth efficiency observed in G2 arrested cells validates that increasing cell size does not automatically result in decreasing growth efficiency even in a model where DNA content does not scale with cell size. Furthermore, these results suggest that G2 growth efficiency is not regulated by dilution of components produced in earlier cell cycle stages.

Finally, using the polyploidy cell data collected using the large-channel SMR, we also analyzed how cell size increase and cell cycle duration scale with cellular hypertrophy and the associated polyploidy. This revealed that with each successive endomitotic cycle the L1210 cells approximately doubled their size independently of the cell size at the start of that cell cycle (Fig. 3G, *top*). Cell cycle duration also remained constant regardless of cell size (Fig. 3G, *bottom).* This suggests that massive cellular hypertrophy and the associated polyploidy do not interfere with the mechanism(s) ensuring that cells double their size during each cell cycle.

In conclusion, increasing cell size does not impose strict transport limitations that would lower growth efficiency in cultured mammalian cells. This conclusion was reached when observing freely proliferating lymphocytes in specific cell cycle stages (Fig. 2C), when examining cells across a vast size range following chemically induced polyploidy (Fig. 3E) and when examining G2 arrested cells *(SI Appendix,* Fig. S3D). Cells may be able to compensate for the increased intracellular distances and decreased surface-to-volume ratio, for example by upregulating the expression of active transporters. Alternatively, the transport limitations may only influence growth under very specific conditions, for example when specific nutrients are in low abundance or when cell size increases above the range normally observed in proliferating mammalian cells.

Our results also show that the previously observed correlation between cell size and growth efficiency in proliferating cultures (1–3) can be explained by cell cycle effects, which have a large influence over cell growth efficiency. While our conclusions rely on data from a single cell line, L1210, the growth profiles of L1210s and adherent cells have been shown to be similar (3), suggesting similar cell cycle dependent growth regulation across multiple cell types. Notably, our results do not exclude growth regulation by dilution effects in G1 stage of cell cycle, nor do our results exclude dilution/concentration effects regulating cell cycle progression or cell metabolism (4, 14–18, 40). In fact, size-dependent dilution effects are likely to be responsible for cell size homeostasis, as our data show that cell size homeostasis is not achieved simply by coupling cell growth efficiency to the absolute size of cells.

Methodologically, we anticipate that the large channel SMRs will have important uses outside this study. The ability to monitor the mass of unlabeled large samples will enable growth (2, 19), drug-response (41) and nutrient uptake (42) studies in various models. These include extremely large single-cells such as adipocytes or megakaryocytes, as well as individual organoids or tumor spheroids, where adherent cell mass accumulation can now be monitored in a preserved 3D microenvironment.

## Materials and Methods

Please see *SI Appendix, Materials and Methods.*

## Acknowledgments

J.H.K. received funding from Samsung scholarship. S.R.M. received funding and support from Koch Institute Frontier Research Program, Koch Institute Support (core) Grant (P30-CA14051), MIT Center for Cancer Precision Medicine, Cancer Systems Biology Consortium funding (CA217377) from the National Cancer Institute. T.P.M. received funding from the Wellcome Trust (110275/Z/15/Z). We would like to thank M. Björklund for useful comments on the manuscript and we would also like to thank M. Stockslager and S. Knudsen for helpful discussions.

## Author contributions

S.R.M. conceived and supervised the large-channel SMR development. T.P.M. conceived and supervised the biological work. L.M., S.O., N.L.C. and R.J.K. designed and set up the large-channel devices, developed their operation interface and optimized their operation. K.R.P. fabricated the large-channel devices. L.M., J.H.K. and T.P.M. carried out the experiments. L.M. and J.H.K. analyzed the data. L.M., J.H.K., S.R.M. and T.P.M. wrote the manuscript with input from all the authors.

## References

1. A. Tzur, R. Kafri, V. S. LeBleu, G. Lahav, M. W. Kirschner, Cell growth and size homeostasis in proliferating animal cells. Science 325, 167–171 (2009).

2. S. Son et al., Direct observation of mammalian cell growth and size regulation. Nat Methods 9, 910–912 (2012).

3. Y. Sung et al., Size homeostasis in adherent cells studied by synthetic phase microscopy. Proc Natl Acad Sci US A 110, 16687–16692 (2013).

4. T. P. Miettinen, M. Bjorklund, Cellular Allometry of Mitochondrial Functionality Establishes the Optimal Cell Size. Dev Cell 39, 370–382 (2016).

5. T. P. Miettinen, M. J. Caldez, P. Kaldis, M. Bjorklund, Cell size control - a mechanism for maintaining fitness and function. Bioessays 39 (2017).

6. S. Soh, M. Banaszak, K. Kandere-Grzybowska, B. A. Grzybowski, Why Cells are Microscopic: A Transport-Time Perspective. J Phys Chem Lett 4, 861–865 (2013).

7. M. Bjorklund, Cell size homeostasis: Metabolic control of growth and cell division. Biochim Biophys Acta Mol Cell Res 1866, 409–417 (2019).

8. K. A. Dill, K. Ghosh, J. D. Schmit, Physical limits of cells and proteomes. Proc Natl Acad Sci U S A 108, 17876–17882 (2011).

9. T. P. Miettinen, M. Bjorklund, Mitochondrial Function and Cell Size: An Allometric Relationship. Trends Cell Biol 27, 393–402 (2017).

10. D. S. Glazier, Beyond the ‘3/4-power law’: variation in the intra-and interspecific scaling of metabolic rate in animals. Biol Rev Camb Philos Soc 80, 611–662 (2005).

11. G. B. West, J. H. Brown, The origin of allometric scaling laws in biology from genomes to ecosystems: towards a quantitative unifying theory of biological structure and organization. J Exp Biol 208, 1575–1592 (2005).

12. S. P. Otto, The evolutionary consequences of polyploidy. Cell 131, 452–462 (2007).

13. B. K. Mable, Ploidy evolution in the yeast Saccharomyces cerevisiae: a test of the nutrient limitation hypothesis. J Evol Biol 14, 157–170 (2001).

14. A. A. Amodeo, J. M. Skotheim, Cell-Size Control. Cold Spring Harb Perspect Biol 8, a019083 (2016).

15. G. E. Neurohr et al., Excessive Cell Growth Causes Cytoplasm Dilution And Contributes to Senescence. Cell 176, 1083–1097 e1018 (2019).

16. K. M. Schmoller, J. J. Turner, M. Koivomagi, J. M. Skotheim, Dilution of the cell cycle inhibitor Whi5 controls budding-yeast cell size. Nature 526, 268–272 (2015).

17. H. Wang, L. B. Carey, Y. Cai, H. Wijnen, B. Futcher, Recruitment of Cln3 cyclin to promoters controls cell cycle entry via histone deacetylase and other targets. PLoS Biol 7, e1000189 (2009).

18. K. M. Schmoller, J. M. Skotheim, The Biosynthetic Basis of Cell Size Control. Trends Cell Biol 25, 793–802 (2015).

19. T. P. Miettinen, J. H. Kang, L. F. Yang, S. R. Manalis, Mammalian cell growth dynamics in mitosis. Elife 8 (2019).

20. M. Shuda et al., CDK1 substitutes for mTOR kinase to activate mitotic cap-dependent protein translation. Proc Natl Acad Sci US A 112, 5875–5882 (2015).

21. T. An, Y. Liu, S. Gourguechon, C. C. Wang, Z. Li, CDK Phosphorylation of Translation Initiation Factors Couples Protein Translation with Cell-Cycle Transition. Cell Rep 25, 3204–3214 e3205 (2018).

22. K. J. Heesom, A. Gampel, H. Mellor, R. M. Denton, Cell cycle-dependent phosphorylation of the translational repressor eIF-4E binding protein-1 (4E-BP1). Curr Biol 11, 1374–1379 (2001).

23. C. Cadart et al., Size control in mammalian cells involves modulation of both growth rate and cell cycle duration. Nat Commun 9 (2018).

24. M. B. Ginzberg et al., Cell size sensing in animal cells coordinates anabolic growth rates and cell cycle progression to maintain cell size uniformity. Elife 7 (2018).

25. D. T. Fox, R. J. Duronio, Endoreplication and polyploidy: insights into development and disease. Development 140, 3–12 (2013).

26. A. C. Lloyd, The regulation of cell size. Cell 154, 1194–1205 (2013).

27. N. J. Ganem, D. Pellman, Limiting the proliferation of polyploid cells. Cell 131, 437–440 (2007).

28. T. A. Zangle, M. A. Teitell, Live-cell mass profiling: an emerging approach in quantitative biophysics. Nat Methods 11, 1221–1228 (2014).

29. C. Cadart et al., Fluorescence eXclusion Measurement of volume in live cells. Methods Cell Biol 139, 103–120 (2017).

30. T. P. Burg et al., Weighing of biomolecules, single cells and single nanoparticles in fluid. Nature 446, 1066–1069 (2007).

31. S. Son et al., Resonant microchannel volume and mass measurements show that suspended cells swell during mitosis. J Cell Biol 211, 757–763 (2015).

32. J. H. Kang et al., Noninvasive monitoring of single-cell mechanics by acoustic scattering. Nat Methods 16,263–269 (2019).

33. C. Cadart, L. Venkova, P. Recho, M. C. Lagomarsino, M. Piel, The physics of cell-size regulation across timescales. Nat Phys 15, 993–1004 (2019).

34. C. O. de Groot et al., A Cell Biologist’s Field Guide to Aurora Kinase Inhibitors. Front Oncol 5, 285 (2015).

35. A. A. Mortlock et al., Discovery, synthesis, and in vivo activity of a new class of pyrazoloquinazolines as selective inhibitors of aurora B kinase. J Med Chem 50, 2213–2224 (2007).

36. M. Tamura et al., Development of specific Rho-kinase inhibitors and their clinical application. Bba-Proteins Proteom 1754, 245–252 (2005).

37. G. B. West, W. H. Woodruff, J. H. Brown, Allometric scaling of metabolic rate from molecules and mitochondria to cells and mammals. Proc Natl Acad Sci U S A 99 Suppl 1, 2473–2478 (2002).

38. L. T. Vassilev et al., Selective small-molecule inhibitor reveals critical mitotic functions of human CDK1. Proc Natl Acad Sci U S A 103, 10660–10665 (2006).

39. A. I. Goranov et al., The rate of cell growth is governed by cell cycle stage. Genes Dev 23, 1408–1422 (2009).

40. A. Litsios et al., Differential scaling between G1 protein production and cell size dynamics promotes commitment to the cell division cycle in budding yeast. Nat Cell Biol 21, 1382-+ (2019).

41. N. Cermak et al., High-throughput measurement of single-cell growth rates using serial microfluidic mass sensor arrays. Nat Biotechnol 34, 1052–1059 (2016).

42. S. Son et al., Cooperative nutrient accumulation sustains growth of mammalian cells. Sci Rep 5 (2015).

