## Supplementary material for "Mass measurements of polyploid lymphocytes reveal that growth is not size limited but depends strongly on cell cycle": Materials and Methods & Supplementary Figures

***SI Appendix* for**

**Mu, Kang, et. al.**

Contains:
Materials and Methods
Supplementary Figures
Supplementary Figure legends

### Materials and Methods

#### *Small-channel SMR details and operation*

Small-channel SMRs were build and operated as detailed in (1, 2). Briefly, the silicon chips that contain the vibrating cantilever were fabricated by CEA-LETI (Grenoble, France). The length of the cantilever used in this study was 350 μm and the buried channel cross section size was 20 × 15 μm (width × height). The small-channel SMR was driven using a piezo-ceramic placed under the device and operated in the second flexural bending mode, with a typical resonance frequency of approximately 1.1 MHz. Piezo-resisters embedded at the base of the cantilever were used to monitor the resonance frequency. The cantilever was actuated using a digital control platform in a direct feedback mode with a typical data rate of 3000 Hz. The SMR device was mounted on a metal clamp with water circulating inside the clamp. The water, and consequently the SMR, was maintained at 37°C by using a temperature-controlled water bath (Julabo). Each input port to the SMR was pressurized with 5% $CO_2$, 21% $O_2$ containing gas (Airgas) to maintain a stable pH (3). The pressure in each input, as well as cantilever actuation, were controlled using a custom-made software (LabVIEW 2016) in order to operate the SMR and control the flow of fluids within the SMR. A typical flow rate within the SMR channels was 2 nL/s, resulting in approximately 200-300 ms transit time through the small-channel SMR cantilever (2). The trapping of a cell within the SMR was obtained by an automated system that adjusts pressures to momentarily stop fluid flow following a resonant frequency change (cell transiting through the cantilever). Approximately 50 s after the fluid flow was stopped, the fluid flow direction was automatically reversed and the measurement of the same cell was repeated. This resulted in a typical measurement interval of 1 min. In addition, the small-channel SMR is equipped with a fluorescence detection setup detailed (1, 4). This consists of a modular Nikon microscope equipped with a Nikon LU plan ELWD 50x/0.55 objective and an 8 mm Voltage Output Type photomultiplier tube (Edmund Optics). Illumination using a Lumencor Spectra X light engine was only activated for 400 ms following cell transit through the cantilever, during which the cell transited through the illumination area and fluorescence signal from the FUCCI cell cycle reporter was detected. We carried out experiments with and without excitation of the FUCCI sensor and did not observe a difference in growth rates. All SMR experiments were carried out by growing the cell within the SMR using the same culture media and chemical perturbations that are detailed below and in figure legends.

#### *Large-channel SMR fabrication and physical dimensions*

Large-channel SMR chips were fabricated on six-inch silicon wafers using a three-level process at the Microsystems Technology Laboratories at MIT. The base device wafer is a double silicon-on-insulator (SOI) substrate (Ultrasil), consisting of a 60 μm silicon layer, a 100 nm buried oxide layer, another 5 μm silicon layer, a 1 um buried oxide layer, and a 500 μm thick silicon handle. Microfluidic channels are etched 60 μm deep in the top silicon layer using deep reactive ion etching (DRIE), with the first buried oxide as etch stop. The base device wafer is then fusion bonded to a device lid wafer, which consists of a standard SOI wafer with 5 μm thick top silicon layer (Ultrasil). The resulting stack is mechanically thinned from the back of the device lid wafer, which yields a structure that contains buried microfluidic channels with 5 μm caps on the top and bottom. To allow fluidic access to the buried channel, an etch through the lid opens the microfluidic channel to the wafer surface. Subsequently, a 70 μm DRIE step defines the cantilever by etching through all of the SOI layers down to the base wafer handle. Microfluidic bypass channels are wet etched into a Pyrex wafer (Bullen) with hydrofluoric acid. Inlet and outlet ports are drilled through the Pyrex wafer with an ultrasonic drilling process. The Pyrex microfluidic wafer is anodically bonded to the top of the device stack. The cantilever is released by DRIE through the silicon from the backside, followed by a brief reactive ion etch to remove the buried oxide. Unlike the small-channel SMRs, these devices were fabricated without piezoresistive readout elements, and the cantilever cavity was left open to atmosphere. See (*SI Appendix*, Fig. S1A) for image of an example large-channel device. All large-channel experiments were carried out on devices with a buried channel cross section size of 60 × 60 µm, and cantilever length of 407 µm.

***Large-channel SMR operation***

The large-channel SMRs were driven using a piezo-ceramic plate underneath the SMR chip. Due to spurious resonances close to the second mode frequency, we operated the devices in the first flexural bending mode. The typical resonance frequency was approximately 420 kHz. In order to overcome position-dependent error at the tip associated with first mode operation, we used flow focusing to confine the transit path of the particle (3). The SMR vibration frequency was measured using an optical lever technique similar to work previously reported (5). The resulting signal is fed into a phase-locked loop at a data rate of 4000 Hz, with a loop bandwidth of 100 Hz (5, 6). Other aspects of the setup, including pressure and temperature control are similar to that of the small-channel SMR. Pressures were controlled to achieve 200-300 ms transit times through the cantilever. In order to improve trap stability, an image-based trapping technique was used to repeatedly measure the cell in the large-channel device. The image-based detection and fluidic control is achieved using a custom-made software in Labview (see *SI Appendix*, Fig. S1B for operating principle). Cells were measured approximately every 30-60 s in each experiment. All SMR experiments were carried out by growing the cell within the SMR using the same culture media and chemical perturbations that are detailed below and in figure legends. Before each experiment, the device was cleaned with 10% bleach followed by a thorough rinse in deionized water. The channel walls are passivated with 1 mg/mL PLL-g-PEG in deionized water for 10 min before flushing the device with measurement media.

***Large-channel SMR measurement error and linearity***

Large-channel SMR calibration was performed using polystyrene beads of known sizes (Duke standard 4000 series purchased from Fisher Scientific) to derive mass sensitivity factors for each device (*SI Appendix*, Fig. S1C-E). The measured frequency changes were scaled by the sensitivity factor to yield buoyant mass. While sensitivity factor was calculated using different sized beads, for consistency all cell data was scaled by calibration factors obtained using 30 µm beads.

The measurement resolution of the large-channel SMRs was determined by repeatedly measuring a single polystyrene bead and calculating the standard deviation of the repeated measurements (Fig. 1B). This reflects error on a single mass measurement (typical measurement interval is approximately 30 s). To assess how our measurement resolution can be improved to capture mass changes that take place over longer time scales, we analyzed the repeated bead measurements by applying moving average filters of various lengths on to the data (Fig. 1C). Overall, the large-channel SMR measurement noise derives from two main sources. First, the SMRs have background frequency noise (~30-50 mHz). Second, as the cell or a particle flows through the cantilever and turns around at the tip of the cantilever, the flow path (i.e. how close to the tip of the cantilever the cell flows) will influence our measurement. To minimize the measurement-to-measurement variability in the flow path, we balance fluid flows so that the cells is maintained close to the outer edge of the cantilever (flow focusing). However, the flow focusing can vary from day-to-day and from cell-to-cell. The large-channel SMR measurement resolution reported in this study reflects error caused by both flow path and frequency background noise. Mass resolution of small and large particles are dominated by frequency noise, whereas intermediate sized particles are dominated by flow focusing noise.

We also evaluated the linearity of the large-channel SMR mass measurements by calculating the mass sensitivity factor for vastly different sized polystyrene beads. Assuming that the different sized polystyrene bead standards are fully comparable in density and shape, these measurements suggested a small deviation from fully linear scaling between buoyant mass and frequency change (*SI Appendix*, Fig. S1E). This small non-linearity could also reflect day-to-day variability in the SMR measurements originating from small changes in flow focusing, room temperature and humidity, solution density etc., or acoustic effects that are specific to hard particles (2). Importantly, the metric of growth efficiency is insensitive to gradual changes in device calibration, as we calculate growth efficiency using small mass intervals accumulated over 15 min or 60 min periods (see *SI Appendix SMR data analysis and statistics*). Although the mass values are affected by calibration, we see approximate doubling of cell mass with each mitosis (Fig. 3G), indicating that this variability is either bead-specific, or the effects are averaged out for cell measurements conducted over multiple days.

#### *Cell culture and chemical treatments*

L1210 FUCCI cells were generated in a previous study (3) and originated from ATCC (Cat#CCL-219). All experiments were carried out using RPMI media that contained 11 mM glucose, 2 mM L-glutamine, 10% FBS (Sigma-Aldrich), 1 mM sodium pyruvate, 20 mM HEPES and antibiotic/antimycotic. All media components were obtained from Thermo Fisher Scientific unless otherwise stated. Cells were grown at 37°C in 5% $CO_2$ and 21% $O_2$ atmosphere. Cells tested negative for mycoplasma.

Barasertib (also known as AZD1152-HQPA) was obtained from Cayman Chemical (Cat#11602), H-1152 was obtained from Sigma-Aldrich (Cat#555550) and RO-3306 was obtained from Cayman Chemical (Cat#15149). We only used fresh stocks (dissolved for <2 weeks) of RO-3306 due to previous observations regarding drug efficacy after longer storage (1). FUCCI expressing L1210 cells displayed higher toxicity following RO-3306 treatment and, consequently, the RO-3306 experiments were carried out using the parental L1210 cell line. All chemicals were dissolved in DMSO and final DMSO concentration was maintained below 0.25%. We did not observe differences in control cell growth with this DMSO concentration. The chemical concentrations used were selected based on cell cycle phenotypes observed in control experiments.

#### *Microscopy*

Microscopy samples were prepared on poly-L-lysine (Sigma-Aldrich, Cat#P8920) coated microscopy slides. L1210 cells were first treated with indicated chemicals and grown under normal culture conditions until one hour prior to fixation when the cells were moved to the microscopy slides. Cells were then washed twice with PBS, fixed with 4% paraformaldehyde for 10 min at RT, washed once with PBS and permeabilized with 0.25% Triton X-1000 in PBS for 10 min at RT. Following one PBS wash the cells were incubated with 5% BSA in PBS for 30 min at RT. Nuclear envelope labeling was carried out o/n at 4°C with Lamin A/C mouse monoclonal antibody conjugated to AlexaFluor 488 (Cell Signaling Technology, Cat#8617) using a 1:50 antibody dilution in PBS supplemented with 5% BSA. The following day, the cells were washed twice with PBS, and membrane labeling was carried out o/n at 4°C with Wheat germ agglutinin conjugated to AlexaFluor 594 (Thermo Fisher Scientific, Cat#W11262) using a 2.5 µg/ml dilution in PBS, as shown before (7). The following day, the cells were washed twice with PBS, and DNA labeling was carried out for 1h at RT with 1:2000 dilution of NuclearMask Blue (Thermo Fisher Scientific, Cat#H10325) in PBS. Following two washes with PBS the cells were mounted using Vectashield mounting medium (Vector Laboratories).

Cells were imaged at RT using DeltaVision wide-field deconvolution microscope with standard filters (DAPI, FITC, Alexa594), a 100x objective and immersion oil with refractive index of 1.516 (Cargille Laboratories). Z-slices were collected with 0.3 µm spacing so that the whole cell volume was covered. Image deconvolution was carried out automatically by the SoftWoRx 7.0.0 software.

Only linear transformations to brightness and contrast were carried out and all images were processed in identical manner. Orthogonal views of representative cells and z-slices were prepared using the 'orthogonal views' function in ImageJ. Location from which the orthogonal view is made from is shown using an orange crosshair in (Fig. 3C). The 3-dimensional z-projections were made using maximum intensity projection function in ImageJ.

The growth and morphology of live L1210 cells was monitored using the IncuCyte live cell analysis imaging system by Sartorius. The images displayed in (*SI Appendix*, Fig. S2B) are an overlay of the phase image and green fluorescence signal image (FUCCI cell cycle sensor) obtained directly from the software without manual corrections. All experiments with Incucyte were carried out by plating cells on a 96-well plate with chemical perturbations in the media and imaging the cells every 1h using 20X objective and 300 ms exposure time for the FITC channel.

***Cell cycle analysis and ploidy estimation***

DNA content was analyzed similarly as before (7). Following indicated chemical treatments, cells were washed with PBS and fixed using ice cold 70% EtOH o/n. The following day, the cells were washed twice with PBS and DNA was stained using FxCycle PI/RNase Staining Solution (Thermo Fisher Scientific, Cat#F10797). After 30 min staining the cells were analyzed using BD Biosciences flow cytometer LSR II HTS with excitation lasers at 561 nm and emission filter at 610/20. Estimations of ploidy levels during SMR measurements were done based on cell mass, as we noticed that cells approximately doubled their mass during every endomitotic cycle.

***SMR data analysis and statistics***

Small-channel SMR frequency data was analyzed using the custom MATLAB code detailed previously (2). Large-channel SMR frequency data was analyzed using similar code, which extracts first mode peaks instead. The resonant frequency changes upon cell transit were converted to buoyant mass using sensitivity factors obtained from polystyrene bead measurements. The resulting buoyant mass measurement and time data were then analyzed for growth rates (pg/h) using the custom MATLAB code published previously (1) with the following modifications:

For small-channel SMR data, mass data were linearly interpolated to have 1000 mass data points for every hour. After interpolation, data was filtered using a moving average of 100 points. Smoothed data was then fitted with a 15 min long linear line, the slope of which is the growth rate, and the fitting was moved 5 min at a time to cover the whole cell cycle. An exception to this is was when growth efficiency was analyzed in a specific cell cycle stages (Fig. 2C), as in these analyses, the line fitting was carried out on a 1 h long segments of raw data without any interpolation.

For large-channel SMR data, the mass data were also interpolated to have 1000 mass data points for every hour and resulting interpolated data was filtered using a moving average of 100 points. Smoothed data was then fitted with a 60 min long linear line, the slope of which is the growth rate, and the fitting was moved 5 min at a time to cover the whole cell cycle.

We assigned the following cell cycle transitions to mass traces: G1/S transition (small-channel SMRs only) was assigned at the point where the mAG-hGeminin signal started increasing; G2/M transition was assigned based on reduced acoustic signal (small-channel SMRs only) and increased mass accumulation rate as detailed before (1, 2); metaphase/anaphase transition was assigned based on loss of mAG-hGeminin signal (small-channel SMRs only), radical drop in acoustic signal (small-channel SMRs only) and minor loss of buoyant mass as detailed before (1, 2); and cytokinesis was assigned as the point where cell mass suddenly reduced by half (abscission of daughter cells). Using these transition points, we excluded mitosis (from G2/M transition to 15 min after metaphase/anaphase transition) from our growth analyses. When analyzing growth efficiency as a function of cell mass in specific points in the cell cycle, we defined newborn G1 as the 60 min period starting 10 min after division, G1/S transition as the 60 min period surrounding the assigned G1/S point, and late G2 as the last 60 min prior to G2/M transition.

To derive the errors in growth efficiency measurements in each specific stage in the cell cycle (Fig. 2C), we utilized the standard deviation of the slope of the fitted linear line ($\delta\ Growth\ rate$) and the standard deviation of a typical mass measurement ($\delta\ mass$) which was determined previously (1). These errors were used to calculate the cumulative error ($\delta$ Growth efficiency) according to the formula:

$$\delta\ \text{Growth efficiency} = |\text{Growth efficiency}|\sqrt{\left(\frac{\delta\ Growth\ rate}{Growth\ rate}\right)^2 + \left(\frac{\delta\ mass}{mass}\right)^2}$$

For all samples, except for RO-3306 treated cells, we only analyzed mass traces that contained one or more mitosis. When cells died while trapped within the SMR, we excluded the part of the data where mass accumulation was zero or negative but included all data prior to the point where mass accumulation stopped (see *SI Appendix*, Fig. S2C for examples). When analyzing control cells using the small-channel SMR, we always monitored the cells for multiple cell cycles to verify that our analysis focused on actively growing and proliferating cells. Partial cell cycles in the control mass traces obtained using small-channel SMRs were not analyzed. No RO-3306 treated mass traces were excluded, but we stopped our data analysis after one doubling of cell mass as long-term drug treatments resulted in cell death. Errors in growth rates and growth efficiencies across the cell cycle (e.g. Figs. 2A and 3E) were plotted by overlaying the individual growth traces and plotting the mean ± standard deviation (representing both technical error and cell-to-cell variability) of the cells at each time point, as previously described (1).

The quantification of cell size dependent growth was carried out using Barasertib treated L1210 cell data from the large-channel SMRs exclusively, in order to minimize influence from drug induced toxicity (between control and drug treated experiments), and from system specific biases, such as differences in shear stress, temperature, light induced toxicity and acoustic effects in the SMR channel (2). The cell size dependent growth was determined based on the slope of a linear line that was fitted to the growth efficiency data spanning five cell cycles (five size doublings) (see *SI Appendix*, Fig. S2F). To minimize influence from cell cycle dependent effects, the fitting was started and stopped at the same point in the cell cycle, by assuming an exact doubling of buoyant mass within each endomitotic cycle. The cell cycle dependent growth efficiency was determined by comparing the maximal and minimal growth efficiency observed within a cell cycle in control cells (excluding mitosis). These cell size and cell cycle dependent growth efficiencies were then compared to the average growth efficiency of control cells during whole cell cycle to derive the % to which cell size and cell cycle influence growth within a typical cell cycle. The error of this % value represents the compound error of average growth rate and that of cell cycle or size dependent growth rate (calculated similarly as the cumulative errors detailed above).

In figures displaying population average growth rates or efficiencies as a function of mass, we only display data points (masses) that have data from three or more independent experiments. All data processing and plotting was done using MATLAB and statistical tests were carried out using Origin Pro 8 software.

***Data availability***

All data is shown in figures and the buoyant mass traces used for the quantitative conclusions of this paper can be found in Dataset S1 attached to this paper.

### Supplementary figures & figure legends

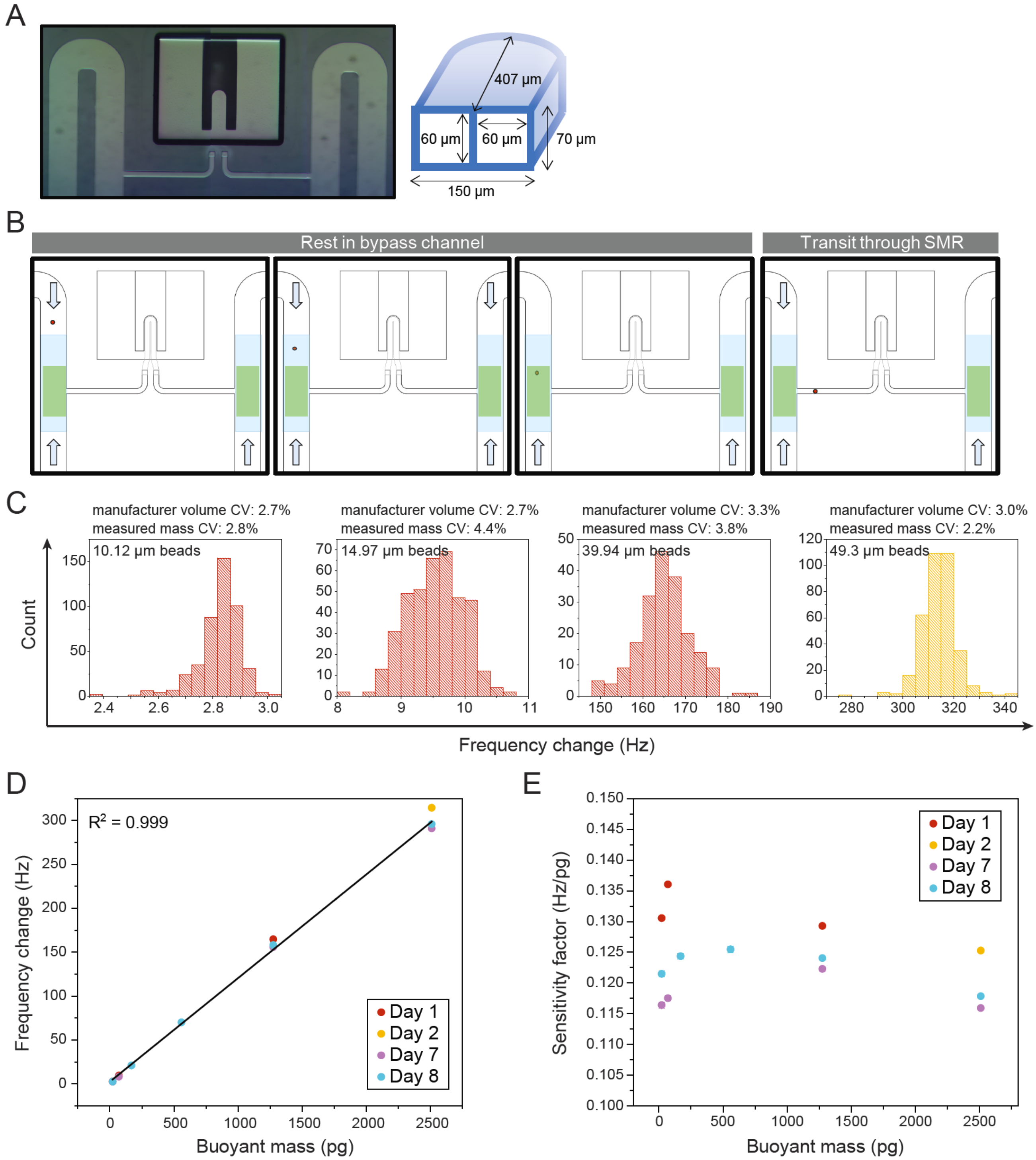


**Fig. S1. Large-channel SMR operates using image-based trapping and expands the analytic measurement range of SMRs.**

**(A)** Microscopy image of a large-channel SMR. Image is reconstructed from three fields of views. A schematic of the large-channel SMR cantilever dimensions are shown on right.

**(B)** Schematic of the image-based hydrodynamic trapping. Four regulators are used to control the pressures at the four ports through which fluid is flown in to the system. Between every measurement the cell remains in the bypass channel for 30-60 s. During this time the image-based hydrodynamic trapping maintains the cell inside the blue shaded region of interest (ROI), but outside the green shaded ROI. If the cell flows out of the ROIs (left side pane), the pressure differences are set to that the cell flows towards the SMR cantilever in the middle. If the cell is inside the blue ROI, but outside the green ROI (middle left pane), then pressures are balanced on all four ports to minimize cell movement. When cell enters green ROI (middle right pane), a minor backwards flow is triggered to prevent the cell from accidentally entering the SMR cantilever. After a defined rest period, the cell pressures are set so that the cell flows through the SMR cantilever to the other side bypass channel (right side pane), where the process of imaging-based hydrodynamic trapping repeats. Blue arrows indicate pressure & fluid flow.

**(C)** Example histograms of 10.12 µm (n = 459 separate beads), 14.97 µm (n = 394 separate beads), 39.94 µm (n = 196 separate beads), and 49.3 µm (n = 351 separate beads) diameter polystyrene bead measurements using the large-channel SMR. Both the measured coefficient of variability (CV) in mass and the manufacturer's reported CV in volume are shown. Data is colored by the date of measurement (same data as Day 1 and Day 2 shown in panels D & E).

**(D)** Large-channel SMR sensitivity scaling obtained by measuring polystyrene beads of different sizes over multiple days. Linear fit and Pearson correlation ($R^2$) are also shown. Frequency change value is the median value of the measured distributions. Error bars are s.e.m. (the error is smaller than each dot). Buoyant mass is calculated based on bead diameter (provided by manufacturer), assuming spherical shape and fixed polystyrene density. Data is colored by the date of measurement. Device is cleaned and passivated between experiments performed on different days (see *SI Appendix Large Channel SMR Operation*), whereas between experiments on the same day the device is flushed thoroughly with buffer.

**(E)** Measured large-channel SMR sensitivity factor as a function of polystyrene bead mass, as measured over multiple days (same data as in panel D). Error bars are s.e.m. (the error is smaller than each dot). See (*SI Appendix Large Channel SMR Measurement Error and Linearity*) for more discussion.

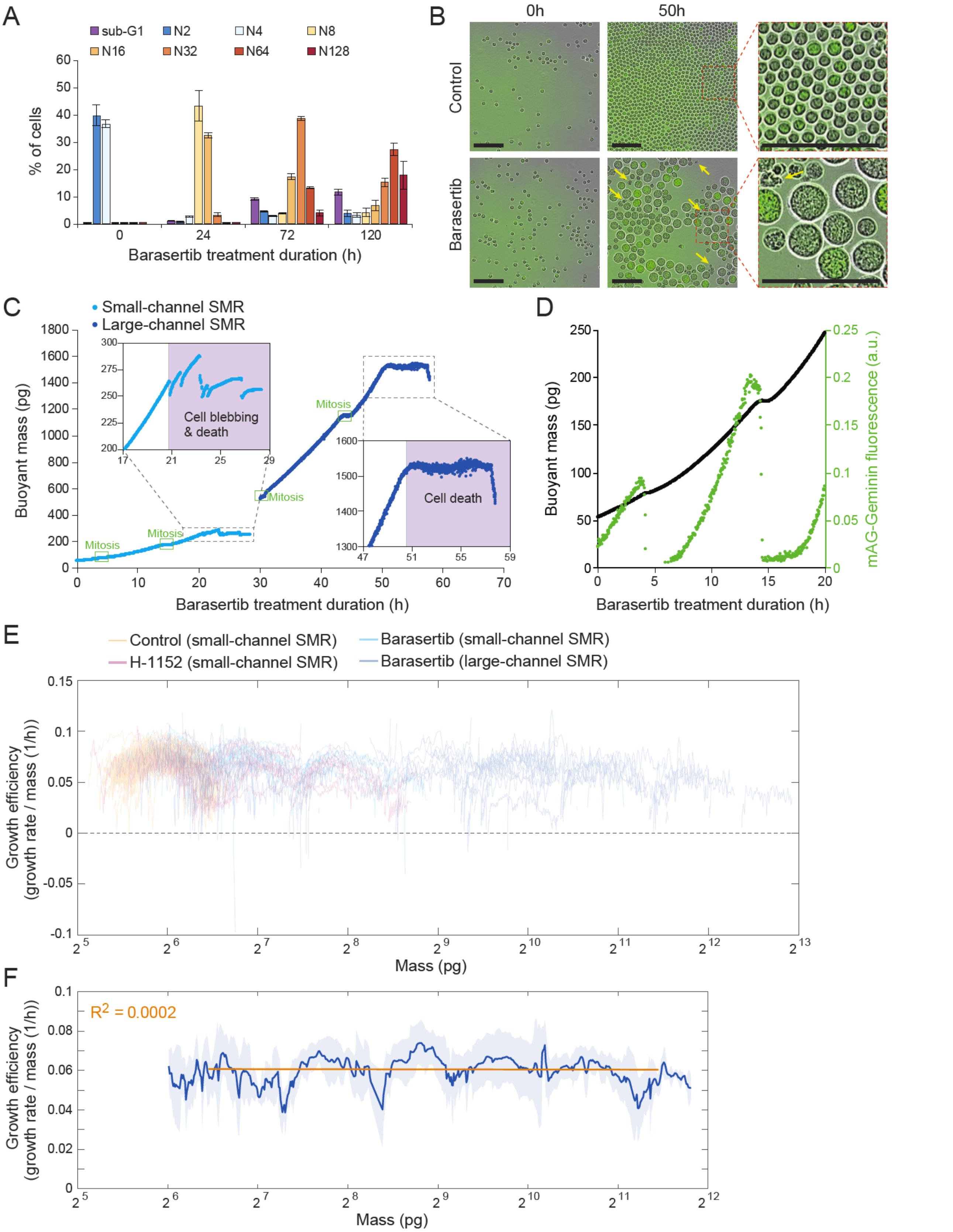


**Fig. S2. Barasertib treatment results in polyploidy, cellular hypertrophy and abrupt cell deaths, without affecting growth efficiency.**

**(A)** Quantification of DNA content (ploidy level) in L1210 FUCCI cells following 50 nM Barasertib treatment (N=3 independent cultures). Note that while cells become polyploid up to 128N, with increasing treatment times also the sub-G1 population increases, indicative of apoptic particles. **(B)** Example phase contrast (grey) and mAG-hGeminin cell cycle reporter (green) overlay images of L1210 FUCCI cells following 50 nM Barasertib treatment (N=2 independent experiments each with 3 independent cultures). Yellow arrows indicate examples of dead/dying cells. Also note the round cell morphology despite a radical volume increase. Scale bars denote 100 μM. **(C)** Two example buoyant mass traces of 50 nM Barasertib treated L1210 FUCCI cells obtained using small (light blue, N=19 independent experiments) and large-channel (dark blue, N=31 independent experiments) SMRs. In these two examples cell growth suddenly stopped, and cells experienced cell blebbing and/or cell death. Once cell growth stopped (light purple areas in the zoom-ins), data was not used to for further analysis. Mitotic events are marked in green and should not be confused with cell death. **(D)** Example buoyant mass (black) and mAG-hGeminin cell cycle reporter (green) trace of a 50 nM Barasertib treated L1210 FUCCI cells obtained using small-channel SMR (N=19 independent experiments). The loss of the mAG-hGeminin signal indicates metaphase-anaphase transition in mitosis and the increase in mAG-hGeminin indicates G1/S transition, thus showing that the cell is undergoing endomitotic cycles. **(E)** The growth efficiency of control (n=72 cells from 9 independent experiments), Barasertib (N=19 independent experiments for small-channel SMR; N=31 independent experiments for large-channel SMR) and H-1152 (N=17 independent experiments) treated L1210 FUCCI cells. Each cell is indicated as an opaque line with gaps between lines representing mitosis, which were not used for the analysis. **(F)** The mean (solid blue line) ± stdev (light blue areas) of growth efficiency as a function of mass for Barasertib treated L1210 FUCCI cells as analyzed by large-channel SMR (N=31 independent experiments). Cell size dependency of growth efficiency was analyzed based on a linear fit (orange line) to a range covering 5 cell size doublings. Pearson correlation ($R^2$) of the fit is displayed in orange.

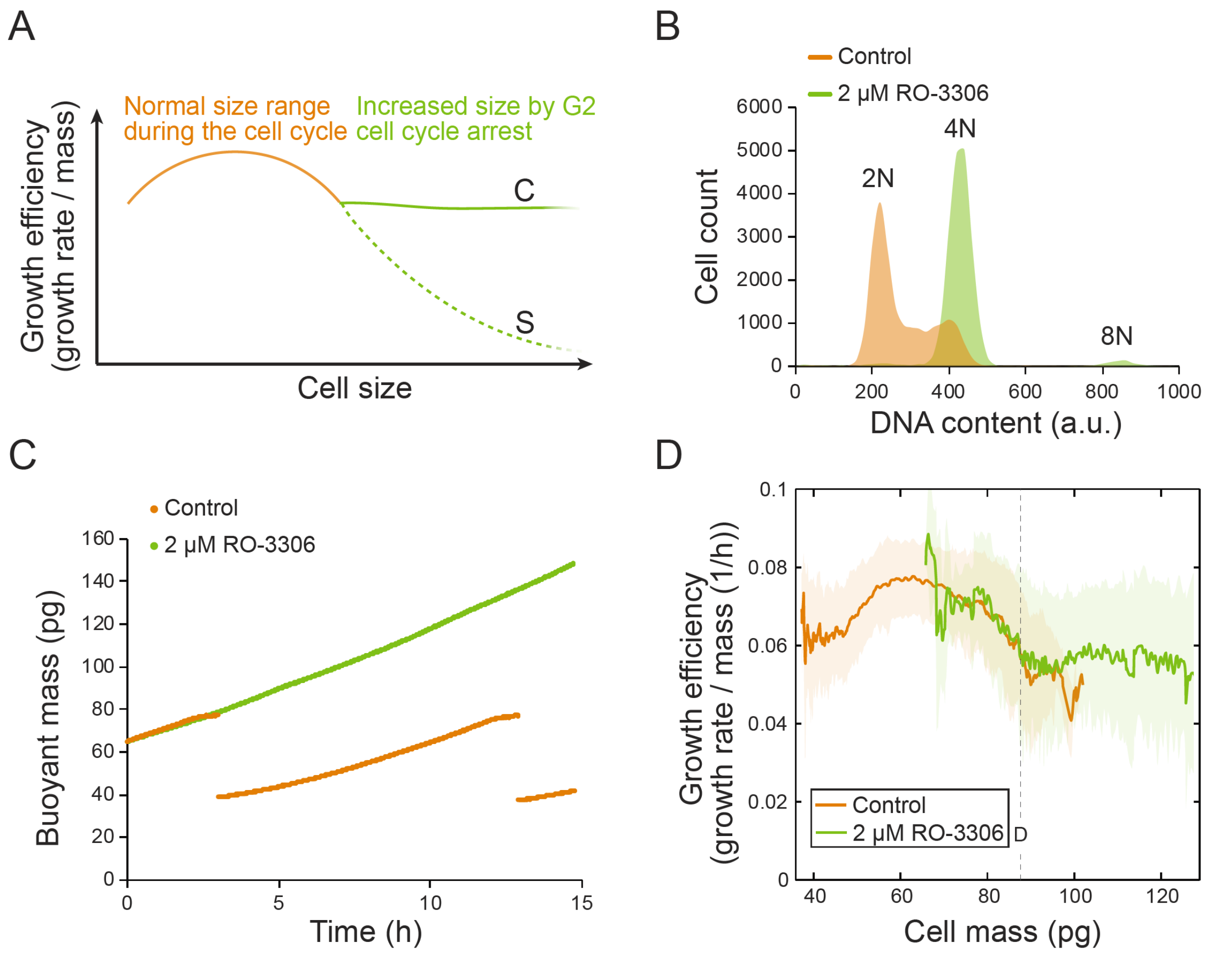


**Fig. S3. G2 cell cycle arrest results in steady growth efficiency.**

**(A)** Hypothesis of cell growth regulation as cells increase size during a G2 cell cycle arrest. In the size-scale observed during a normal cell cycle (orange) cells display non-linear size scaling of growth rates. When cell cycle is arrested in G2, the cell size-dependent (S, dashed green line) and cell cycle-dependent (C, solid green line) growth regulations should result in different growth behaviors. **(B)** Example DNA histogram of control (orange) and 24h RO-3306 treated (green) L1210 cells (N=3 independent cultures). RO-3306 results in a G2 arrest and most cells do not undergo endoreplication cycles. Longer treatments resulted in sub-G1 population increasing, indicative of cell death, which is consistent with previous reports of CDK1 inhibition in vitro (8). **(C)** Example buoyant mass trace of a control (orange) and RO-3306 treated (green) L1210 cell obtained using small-channel SMR (N=9 independent experiments for control; N=12 independent experiments for RO-3306). **(D)** The growth efficiency of control (orange) and RO-3306 treated (green) L1210 cells (N=9 independent experiments for control; N=12 independent experiments for RO-3306). For all experiments with RO-3306, chemical treatment times were under 24h. Solid lines indicate the average growth efficiency and shaded areas display ± stdev. Dashed vertical line indicates the typical division size (D) of control cells. As the G2 arrested cells grow larger than the normal division size, growth efficiency remains constant.

### Supplementary references

1. T. P. Miettinen, J. H. Kang, L. F. Yang, S. R. Manalis, Mammalian cell growth dynamics in mitosis. *Elife* **8** (2019).
2. J. H. Kang *et al.*, Noninvasive monitoring of single-cell mechanics by acoustic scattering. *Nat Methods* **16**, 263-269 (2019).
3. S. Son *et al.*, Direct observation of mammalian cell growth and size regulation. *Nat Methods* **9**, 910-912 (2012).
4. J. H. Kang *et al.*, Time-dynamics of mitochondrial membrane potential reveal an inhibition of ATP synthesis in mitosis. *BioRxiv* 10.1101/772244 (*2019*).
5. N. Cermak *et al.*, High-throughput measurement of single-cell growth rates using serial microfluidic mass sensor arrays. *Nat Biotechnol* **34**, 1052-1059 (2016).
6. S. Olcum, N. Cermak, S. C. Wasserman, S. R. Manalis, High-speed multiple-mode mass-sensing resolves dynamic nanoscale mass distributions. *Nat Commun* **6**, 7070 (2015).
7. T. P. Miettinen, M. Bjorklund, Mevalonate Pathway Regulates Cell Size Homeostasis and Proteostasis through Autophagy. *Cell Rep* **13**, 2610-2620 (2015).
8. M. Bjorklund *et al.*, Identification of pathways regulating cell size and cell-cycle progression by RNAi. *Nature* **439**, 1009-1013 (2006).
